# Bardet-Biedl Syndrome 1 Mutations Differentially Impact BBSome Integrity and its Function in Ciliary Trafficking

**DOI:** 10.64898/2026.01.23.701255

**Authors:** Kristyna Maskova, Hana Hajsmanova, Sofiia Bykova, Sindija Smite, Avishek Prasai, Daniel Rozbesky, Martina Huranova

## Abstract

Bardet-Biedl Syndrome (BBS) is a pleiotropic ciliopathy marked by retinal degeneration, obesity, polydactyly, renal and reproductive anomalies, and cognitive impairment. *BBS1*, the most frequently mutated gene in BBS, encodes a key component of the BBSome complex, which is essential for ciliary membrane trafficking.

Although BBS1 is known to be essential for proper BBSome function, the effects of disease-associated BBS1 variants on its activity remain incompletely understood. In this study, we examined how patient-derived BBS1 mutations affect BBSome integrity and its role in cargo transport within primary cilia. Our results show that particular BBS1 mutations interfere with distinct stages of BBSome assembly and trafficking. While M390R disrupts initial pre-BBSome assembly at pericentriolar satellites, E224K impairs both the maturation of the pre-BBSome into the BBSome and its movement from pericentriolar satellites to the cilium. In contrast, the R160Q variant preserves BBSome assembly and permits its localization to cilia. It specifically weakens the BBSome-GPCR interaction mediated by TOM1L2, resulting in defective GPR161 export and increased ciliary IFT turnover.

Overall, our study establishes a mechanistic framework linking specific BBS1 mutations to distinct defects in BBSome assembly and function. This framework defines functional classes of BBS1 variants and provides deeper insight into the molecular mechanism and severity of Bardet-Biedl Syndrome.

## Introduction

Bardet-Biedl syndrome (BBS) (OMIM 209900) is the most common non-lethal multisystemic genetic disorder caused by dysfunction of the primary cilium, a microtubule based sensory organelle. BBS presents with a broad spectrum of clinical features, including retinal degeneration, polydactyly, obesity, renal anomalies, cognitive impairment, and hypogonadism ^1,2^. To date, at least 26 genes have been linked to BBS, eight of which encode subunits of the BBSome complex: BBS1, BBS2, BBS4, BBS5, BBS7, BBS8, BBS9, and BBS18 ^3–5^. The BBSome exports ciliary cargo via interactions with intraflagellar transport machinery (IFT) and TOM1L2, an adapter for ubiquitinated cargoes ^6–11^. The BBSome regulates multiple signaling pathways, such as Sonic Hedgehog ^12–14^, WNT ^15–17^, photoreceptor signaling ^18,19^, and neuronal signaling mediated by G protein-coupled receptors (GPCRs) ^20,21^.

BBS1, a key subunit of the BBSome, is the most commonly mutated gene in BBS, accounting for approximately 28% of cases ^4,22^. Structural and biochemical studies have demonstrated that the BBS1 plays a central role in BBSome assembly and function. It directly interacts with multiple BBSome subunits, contributes to the formation of the cargo-binding sites, binds the small GTPase ARL6/BBS3 to facilitate BBSome activation, membrane recruitment and translocation to cilia ^5–7,23–26^.

BBS1 mutations are typically associated with a milder clinical form of BBS than mutations in some other BBSome encoding genes ^4^, with the symptom frequency and severity varying depending on the specific mutation involved; substitution mutations also tend to have milder symptoms than larger gene aberrations ^4,27–30^. The most common BBS1 variant is the M390R missense mutation, a major pathogenic allele in BBS ^4,27^, likely disrupting the BBS1 stability and interaction with ARL6 and BBS4 ^23,31–33^. The E224K and R160Q mutations are linked to milder clinical symptoms that, based on clinical presentation alone, may not always fulfil the diagnostic criteria for BBS ^1,2,28–30^. While additional BBS1 mutations have been identified ^34,35^, their pathogenic effects are primarily inferred from structural models of the BBSome ^25,26^.

In this study, we found that the BBS1 mutations M390R, E224K, and R160Q can be divided into distinct functional classes: those that disrupt BBSome assembly and its translocation to cilia, and those that permit BBSome assembly but fail to facilitate cargo retrieval by connecting cargo-bound TOM1L2 to IFT. These results show unique pathological mechanisms for individual genetic mutations underlying BBS.

## Results

### Pathogenic BBS1 patient mutations selectively disrupt the stability of the BBSome subunits

Mutations in the BBS1 subunit of the BBSome are associated with comparatively milder Bardet– Biedl syndrome (BBS) phenotypes than mutations in core BBSome subunits BBS2, BBS7, and BBS9 ^4^. Our work, along with that of others, has shown that loss of BBS1 results in accumulation of the pre-BBSome at pericentriolar satellites, whereas loss of other BBSome subunits disrupts pre-BBSome nucleation at these sites ^5,24^. To investigate the pathogenic mechanisms by which BBS1 mutations contribute to BBS, we selected three variants with varying clinical penetrance: M390R, E224K, and R160Q (Fig. 1A). These mutations map to distinct structural regions of BBS1, enabling analysis of region-specific effects on BBSome assembly and function (Fig. 1A-B) ^7,25^. The M390R mutation lies in a buried hydrophobic cluster and introducing a charged arginine at this position destabilizes the local β-propeller packing, consistent with earlier reports showing loss of ARL6–GTP binding, and diminished BBS4 interaction ^23,33^. The E224K mutation lies at the BBS1–BBS4 interface and introduces a charge inversion that is likely to weaken the interaction by disrupting local electrostatic complementarity. The R160Q variant could disrupt an intramolecular salt bridge within the flexible 4α-helical region that contacts BBS4, increasing local flexibility and potentially loosening this interface.

**Figure 1.**
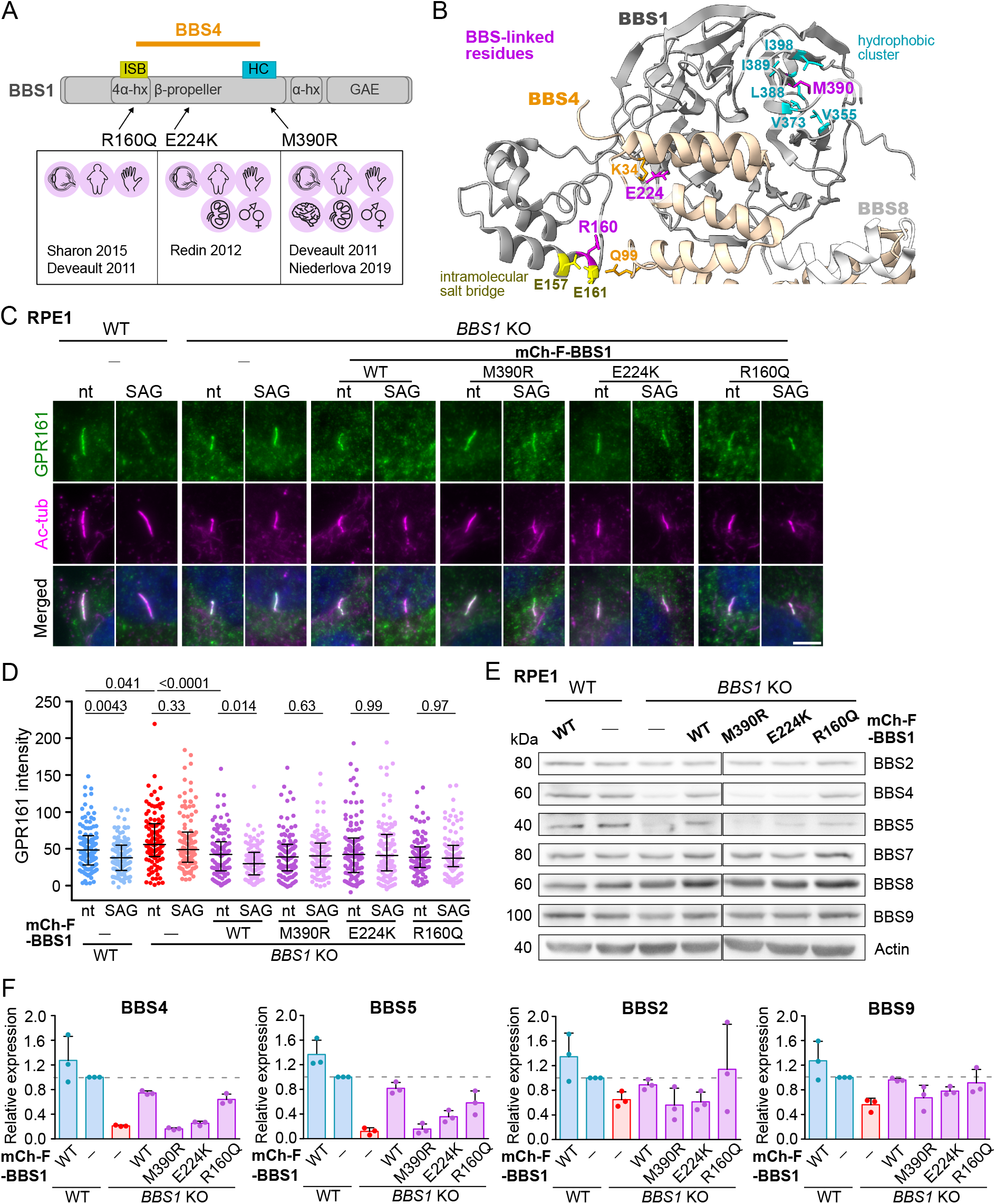
BBS linked BBS1 variants differentially alter expression of the BBSome subunits. (A) Schematic representation of the BBS1 subunit of the BBSome showing its domain organization and the locations of BBS–associated mutations of varying clinical severity (indicated by organ-specific icons and respective studies). Based on structural analyses, the mutations map to the BBS1– BBS4 interaction interface (orange line). ISB, intramolecular salt bridge; HC, hydrophobic cluster; hx, α-helix. (B) Ribbon representation of the BBS1–BBS4 interaction interface from the cryo-EM structure of the human BBSome ^7^, shown together with a portion of BBS8 (white). The N-terminal β-propeller of BBS1 is shown in grey and the N-terminal TPR domain of BBS4 in orange; BBS-linked residues in BBS1 are highlighted in magenta. M390 of BBS1 is located in blade 7 of the β-propeller and is buried within a hydrophobic cluster formed by V355, V373, L388, I389, and I398 (cyan), distal from the BBS1–BBS4 interface. E224 of BBS1 lies at the BBS1–BBS4 interface in blade 4 of the β-propeller and is positioned near K34 of BBS4 (orange). R160 is located within the 4α-helical insert of the BBS1 β-propeller, where it forms an intramolecular salt bridge with E161 and E157 (yellow); this helical insert contacts the TPR domain of BBS4, with the R160 backbone positioned near Q99 of BBS4 (orange). Highlighted residues are shown as sticks. Representative micrographs (C) and intensity quantification (D) of GPR161 in cilia in the parental and BBS1 variants expressing WT and *BBS1* KO cell lines starved for 24 h upon treatment with 200 nM SAG for 2h. Antibody against acetylated tubulin (Ac-tub) was used to stain cilia. Nucleus was stained with 4′,6-diamidino-2-phenylindole (DAPI). Scale bar, 5µm. Analysis was carried out using the Fiji ImageJ. Medians with interquartile range from three independent experiments of (n = 110-140 cilia). Statistical significance was calculated using two-tailed Mann-Whitney test. (E) Expression levels of endogenous BBSome subunits in parental and BBS1 variants expressing WT and *BBS1* KO RPE1 cell lines. Equal protein amounts were loaded into each lane. Actin is used as the loading control. Representative blots out of three independent experiments are shown. (F) Bar graphs depicting the relative expression levels of the endogenous BBSome subunits in parental and BBS1 variants expressing WT and *BBS1* KO RPE1 cell lines. Protein amounts were quantified using the Fiji ImageJ. Protein expression was first normalized to actin and then to levels in parental WT cell line. Average and SD of three independent experiments is shown.

To characterize the role of these mutations in the BBSome biology, we generated BBS1 variants tagged at the N-terminus with mCherry and FLAG (mCh-F-BBS1). These constructs were introduced into RPE1 *BBS1* knockout (KO) cells ^5^. Since the M390R and R160Q mutations have been associated with protein degradation in BBS patients ^31,32^, we first examined their protein expression in our model. We sorted cells double positive for mCherry and Thy1.1, a marker encoded in a bicistronic mCh-F-BBS1∼IRES-Thy1.1 transcript in the pMSCV vector (Fig. S1A). Correlation analysis of mCherry and Thy1.1 levels indicated that the mCh-F-BBS1 variants are stable, with M390R, E224K, and R160Q showing even higher stability than WT (Fig. S1A-B). Analysis of BBS1 levels showed that the variants were expressed at levels approximately 40-fold higher than endogenous BBS1 (Fig. S1C-D). This allowed us to study their effects on BBSome assembly and function without interference from potential variant instability or degradation.

Next, we examined how these variants affect BBSome function in cilia. We assessed BBSome-mediated GPCR retrieval during signaling by focusing on the canonical BBSome cargo GPR161, an inhibitory GPCR in the Hedgehog (HH) pathway ^36^. A key feature of HH pathway activation is the BBSome-dependent removal of GPR161 from cilia ^10,13,24,37^, which occured in WT cells and *BBS1* KO cells complemented with mCh-F-BBS1 WT (Fig. 1C-D). In contrast, all three BBS1 variants - M390R, E224K and R160Q - failed to restore GPR161 export in the *BBS1* KO background, demonstrating their sl of function (Fig. 1C–D).

Our previous study demonstrated that deletion of BBS1 reduces the expression of several BBSome subunits—most prominently BBS4 and BBS5, and to a lesser extent BBS2, BBS7, BBS8, and BBS9—owing to impaired BBSome assembly ^5^. Expression of mCh-F-BBS1 WT restored BBSome subunit levels in the *BBS1* KO background (Fig. 1E-F, S1E). In contrast, the M390R and E224K variants failed to rescue the reduced subunit expression, whereas the R160Q variant unexpectedly restored expression nearly to WT levels (Fig. 1E-F, S1E). Together, these results indicate that pathogenic BBS1 mutations differentially influence the stability of BBSome subunits.

### BBS1 variants differentially disrupt BBSome assembly and ciliary localization

Fully assembled BBSome localizes to cilia, which is essential for its function ^5,6^. Since BBS1 completes BBSome assembly and facilitates its BBS3-dependent translocation to cilia ^5,6,23^, we examined BBS9 localization in cells expressing mCh-F-BBS1 variants by immunofluorescence (Fig. 2A-C, S2A-B). Expression of either the mCh-F-BBS1 WT or R160Q variant restored BBS9 localization to cilia at comparable frequencies and with a comparable distribution along the cilia, indicating that the R106Q mutation does not impair BBSome entry into the cilium. In contrast, neither BBS9 nor BBS1 localized to cilia in cells expressing the mCh-F-BBS1 M390R or E224K variants (Fig. 2A-B, S2A).

**Figure 2.**
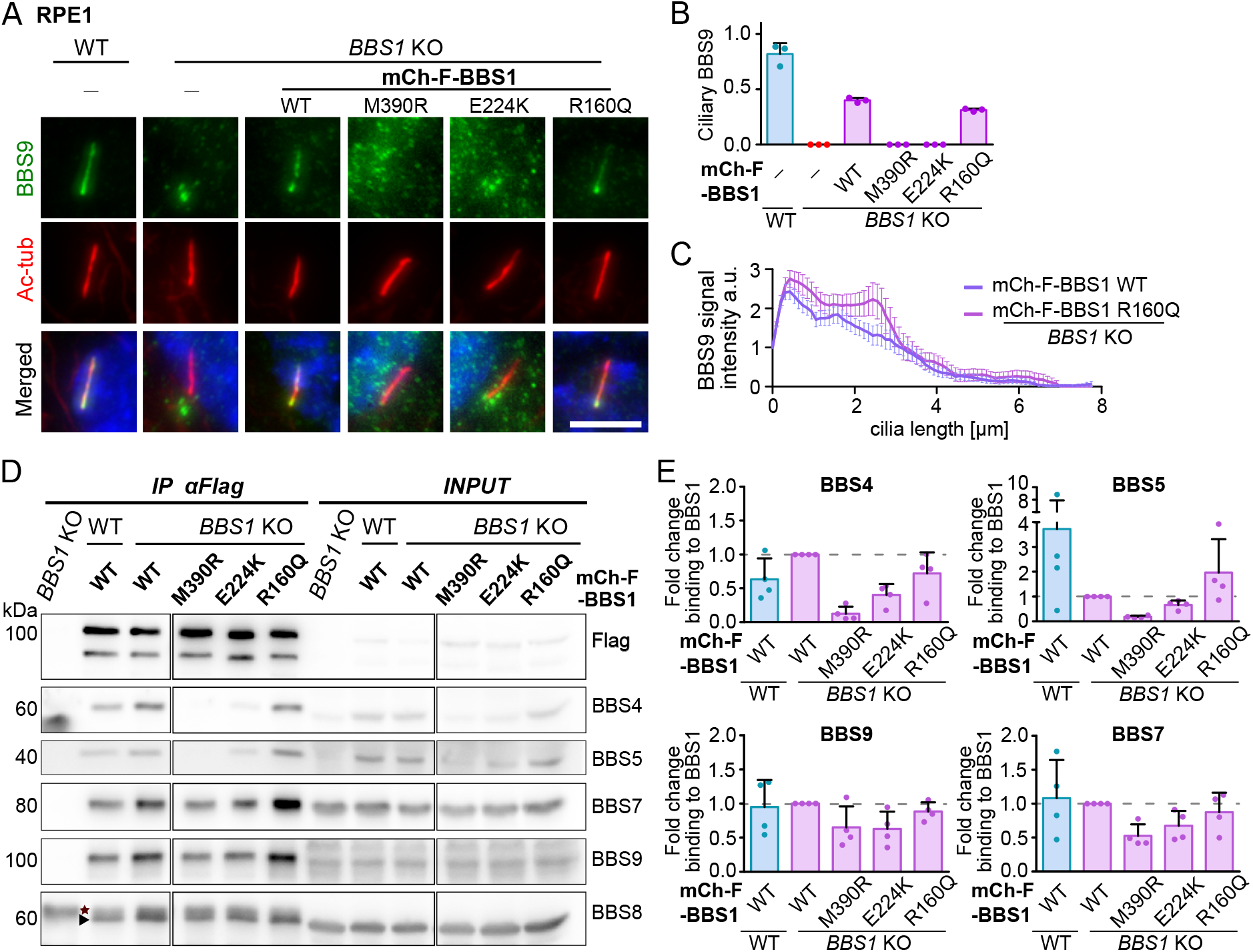
BBS1 variants differentially impact BBSome assembly and translocation to cilia. Representative micrographs depicting BBS9 localisation (A) and quantification of ciliary BBS9 (B) in the parental and BBS1 variants expressing WT and *BBS1* KO RPE1 cell lines upon 24 h serum starvation. Antibody against acetylated tubulin (Ac-tub) was used to stain the primary cilia. Nucleus was stained with 4′,6-diamidino-2-phenylindole (DAPI). Scale bar, 5µm. Analysis was carried out using the Fiji ImageJ. Mean and SD from three independent experiments (n = 160-230 cilia). (C) Distribution of BBS9 signal along the axoneme in *BBS1* KO RPE1 cells expressing BBS1 WT and R160Q variants in (A). Quantification was performed using the Fiji ImageJ. The plot shows mean and SD from n = 41 cilia for WT and n = 35 cilia for the R160Q variant. (D) Co-immunoprecipitation of the BBSome subunits from mCherry-FLAG tagged BBS1 variants expressing WT and *BBS1* KO RPE1 cell lines using anti-FLAG antibodies after 24 h serum starvation. Representative immunoblot out of three experiments is shown. (E) Quantification of the co-precipitated BBSome subunits in (D) was performed using the Fiji ImageJ software. Amount of the BBSome subunits was normalized to the levels of the mCherry-FLAG tagged BBS1 WT in *BBS1* KO RPE1 cell line detected on the respective membrane. Mean with SD of three experiments is shown.

To assess the formation of BBSome subcomplexes, we performed FLAG-based immunoprecipitation assays (Fig. 2D-E, S2C). The mCh-F-BBS1 R160Q variant co-precipitated BBSome subunits to comparable levels to the BBS1 WT. This suggests that the R160Q mutation does not block BBS1 integration into the BBSome or its targeting to cilia and may underlie BBS pathology through BBSome dysfunction within cilia. In contrast, the mCh-F-BBS1 M390R and E224K variants showed markedly reduced association with the BBS4 and BBS5 subunits and retained substantial interactions with the BBSome core components (BBS7 and BBS9), as well as with the BBS8 subunit (Fig. 2D-E, S2C).

Collectively, these findings suggest that BBS-linked mutations in BBS1 can perturb specific steps of BBSome assembly and function without necessarily disrupting completely the BBSome assembly.

### E224K and M390R mutations in BBS1 disturb pre-BBSome formation at pericentriolar satellites

At pericentriolar satellites, BBS4 together with BBS9 recruit the BBSome subunits to initiate pre-BBSome assembly, which is completed by the addition of BBS1 at the basal body ^5^. Our data show that the M390R and E224K mutations in BBS1 alter its interaction with BBS4 to different extent (Fig. 2D-E). To assess the impact of BBS1 variants on BBS4-driven pre-BBSome assembly at satellites, we analysed BBS9 co-localization with PCM-1, a satellite marker (Fig. 3A). In the presence of R160Q variant, BBS9 was devoid from satellites to a similar extent as with the WT variant, in both *BBS1* KO and WT backgrounds (Fig. 3B), consistent with the BBSome`s ability to translocate to cilia (Fig. 2A). Interestingly, the E224K variant promoted BBS9 accumulation at satellites to levels comparable to those in the *BBS1* KO condition, indicating formation of the pre-BBSome (Fig. 3B). By contrast, in the presence of M390R variant, BBS9 localized to satellites to an even lesser extent compared to WT and R160Q conditions (Fig. 3A-B). Since the M390R variant associates with BBS9, BBS7 and BBS8, but not with BBS4 and BBS5 subunits (Fig. 2A, E, S2C), these data suggest that this subcomplex may be depleted from satellites into the cytoplasm.

**Figure 3.**
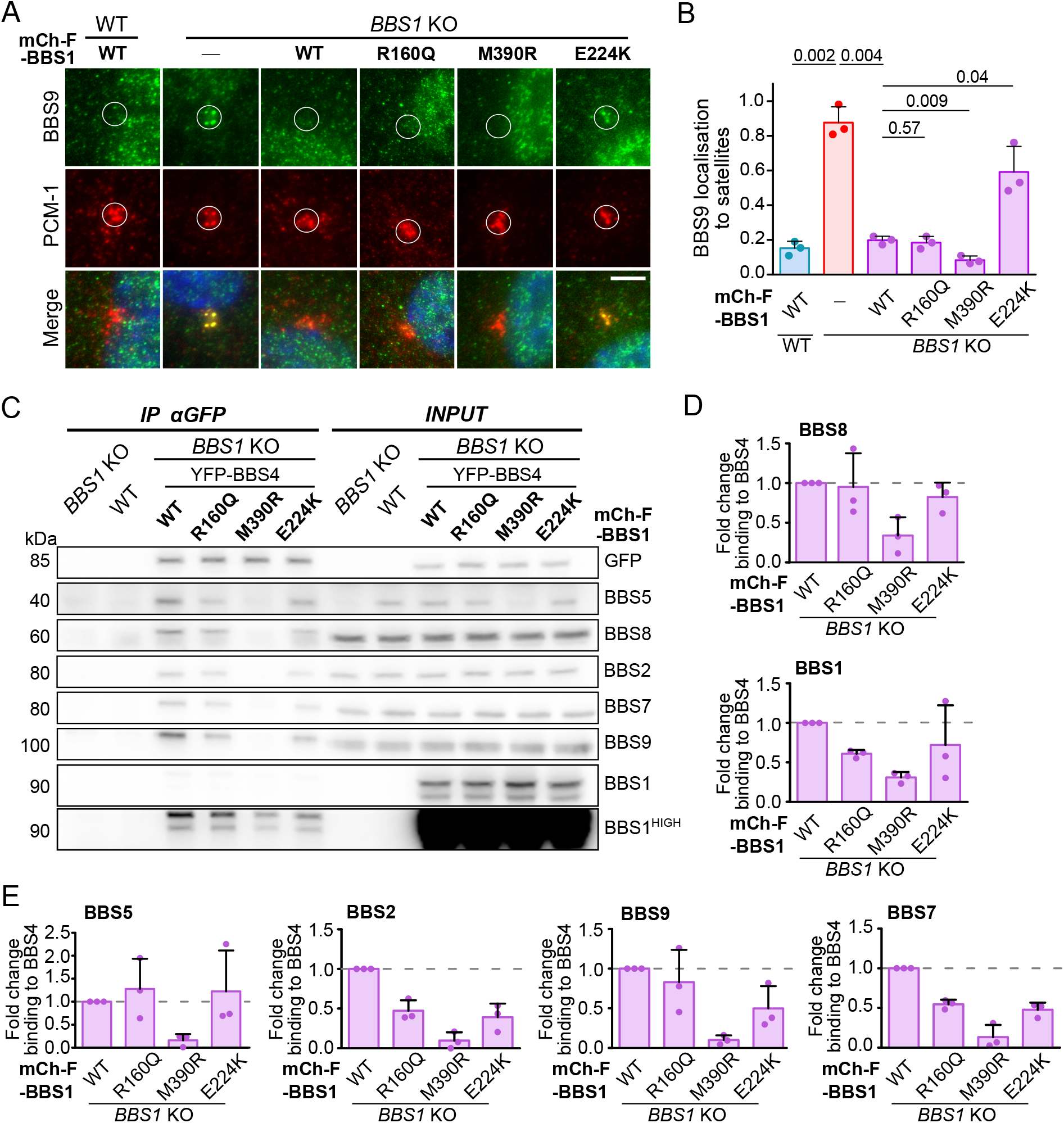
BBS1 E224K and M390R alter the pre-BBSome assembly at pericentriolar satellites. Representative micrographs (A) and quantification (B) showing localisation of BBS9 in pericentriolar satellites in the parental and BBS1 variants expressing WT and *BBS1* KO RPE1 cell lines. Antibody against PCM-1 was used to stain the satellites. Nucleus was stained with 4′,6-diamidino-2- phenylindole (DAPI). Scale bar, 5µm. Analysis was carried out using the Fiji ImageJ software. Mean and SD from three independent experiments (n = 220-340 satellites). Statistical significance was calculated using two-tailed paired t-test. (C) Co-immunoprecipitation of the BBSome subunits from YFP-BBS4 and BBS1 variants expressing *BBS1* KO RPE1 cell lines using anti-GFP antibodies after 24 h serum starvation. Representative immunoblot out of three experiments is shown. Quantification of the co-precipitated BBS8 and BBS1 (D) and BBS5 and BBSome core subunits (E) in (C) was performed using the Fiji ImageJ. Amount of the BBSome subunits was normalized to the YFP-BBS4 levels in the mCherry-FLAG tagged BBS1 WT expressing *BBS1* KO RPE1 cell line, as detected on the respective membrane. Mean with SD of three experiments is shown.

To explore this further, we expressed YFP-tagged BBS4 in the cells and used YFP-based immunoprecipitation to pull out the complexes (Fig. 3C-E, S3A). Because YFP-BBS4 mainly localizes to the centrosome-associated satellites in all BBS1-variant cell lines (Fig. S3A), the complexes we isolated reflected also interactions happening at those sites. The WT, R160Q and E224K variants supported pre-BBSome/BBSome complex formation, with BBS4 efficiently binding BBS8, BBS1, BBS5 and substantial amounts of the core BBSome components (Fig. 3D-E). The E224K mutation enabled incorporation of BBS1 into the pre-BBSome, yet fraction of the assembled E224K-BBSome retained at satellites (Fig. 3A, B, D). We previously showed that, BBS4 dissociates faster from satellites in the absence of BBS1, reflecting pre-BBSome instability ^5^. Using FRAP, we found that BBS4 dissociation in the presence of BBS1 E224K is comparable to WT, indicating that the E224K-BBSome is neither destabilized nor sequestered at satellites (Supp. Fig. 3B–D). These results show that the BBS1 E224K mutation allows BBSome assembly but specifically impairs its translocation to cilia (Fig. 2A-B), likely due to compromised complex integrity (Fig. 1B).

In the presence of the BBS1 M390R variant, BBS4 bound only minimal amounts of BBS1 and BBS8, and virtually no BBS5 or core BBSome components BBS2, BBS7, and BBS9 (Fig. 3D–E). Despite this, the BBSome core still associated directly with M390R (Fig. 2D–E), while BBS9 levels at satellites remained low (Fig. 3B), suggesting that M390R sequesters the core away from BBS4 and satellites, preventing pre-BBSome nucleation. In contrast, BBS1 knockout cells showed clear enrichment of BBS9 and other subunits at satellites (Fig. 3A-B)^5^, and modestly increased BBS4 association with BBS9 and BBS7 (Supp. Fig. 3E–F).

Altogether, these data suggest that E224K behaves as a hypomorphic allele, with a milder effects than *BBS1* KO, likely due to a fraction of partially functional BBSome at the satellites. In contrast, M390R more closely resembles *BBS1* KO, as it interferes with pre-BBSome formation, albeit via a distinct mechanism.

### BBS1 R160Q disrupts TOM1L2-GPCR coupling to retrograde IFT

We found that BBS-linked BBS1 mutations perturb BBSome function through distinct mechanisms: they disrupt BBSome assembly, transport to cilia, or function within cilia. To investigate why the BBSome-BBS1 R160Q complex cannot export GPR161, we examined the ciliary localisation and abundace of TOM1L2, an adaptor for ubiquitinated cargoes targeted for BBSome-dependent exit (Fig. 4A-B). As reported previously, TOM1L2 accumulates in cilia of cells lacking the BBSome ^11^. Consistent with this, we observed increased abundace of ciliary TOM1L2 in cells expressing the BBS1 M390R and E224K variants, in which the BBSome fails to form (Fig. 2A, D). Notably, TOM1L2 also accumulated in cells with the BBSome-BBS1 R160Q complex, indicating this variant impairs BBSome-mediated export of the TOM1L2-marked cargoes (Fig 4A-B).

**Figure 4.**
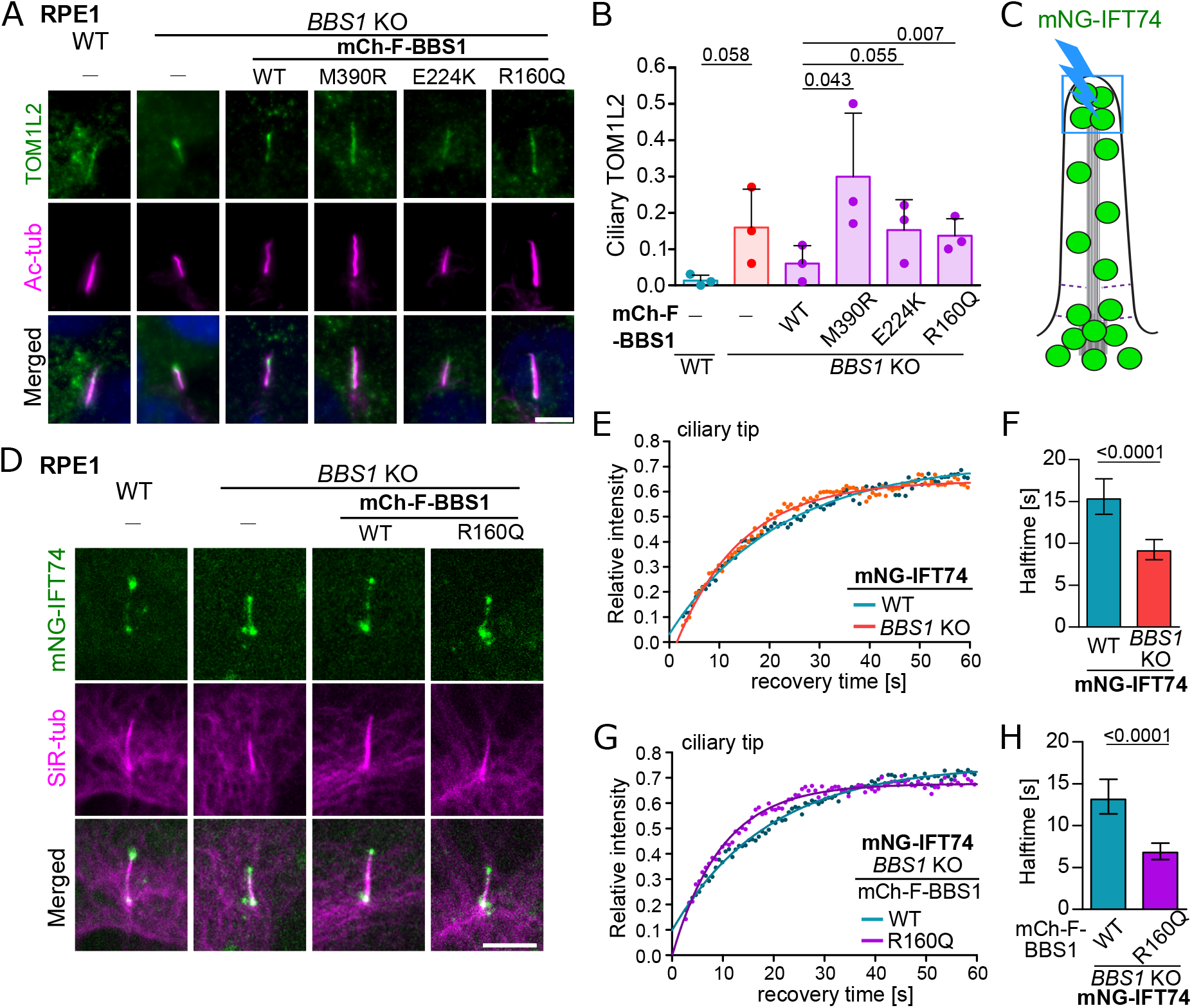
BBS1 R160Q disrupts cargo binding and retrieval of BBSome dependent GPCRs. Representative micrographs (A) and frequency quantification (B) of TOM1/2L in cilia in the parental and BBS1 variants expressing WT and *BBS1* KO RPE1 cell lines starved for 24 h. Antibody against acetylated tubulin (Ac-tub) was used to stain cilia. Nucleus was stained with 4′,6-diamidino-2- phenylindole (DAPI). Scale bar, 5µm. Analysis was carried out using the Fiji ImageJ. Means with SD from three independent experiments of (n = 290-350 cilia). Statistical significance was calculated using one-tailed paired t-test. (E) Schematic representation of the FRAP experiment in the cilia localised IFT74-mNeonGreen (green circles). Ciliary tip (blue box) was bleached for 0.65 s using a 20 mW 488 nm solid-state laser at 100% intensity (blue lightning bolt). (F) Representative micrographs from the FRAP experiments depicting localisation of IFT74- mNeonGreen in the parental and BBS1 WT and R160Q variants expressing WT and *BBS1* KO RPE1 cell lines. Ciliary axonemes and microtubules were stained with 1μM SiR-tubulin. Scale bar, 5µm. (E, G) FRAP analysis of the mNG-tagged IFT74 in the parental and BBS1 WT and R160Q variants expressing WT and *BBS1* KO RPE1 cell lines upon 48 h serum starvation. Recovery curves were fitted at once using one phase association fit. Means of (n = 45 - 50) measurements from four independent experiments are shown. (F, H) Bar graphs depicting recovery halftimes (s) derived from curve fits in (E, G). Error bars represent the 95% confidence interval and p-value is shown.

Since the BBS1 R160Q variant allows BBSome assembly but not cargo export, we further examined whether it disturbs TOM1L2-cargo loading onto IFT. Studies in olfactory neurons, *C. elegans*, and *Chlamydomonas* showed that the BBSome associates with the IFT machinery and regulates IFT dynamics and turnaround at the ciliary tip ^9,38–40^, where anterograde trains are remodelled to initiate retrograde transport ^41^. First, we investigated how the BBSome regulates mammalian IFT dynamics by measuring dwell time of IFT74–mNeonGreen (IFT74–mNG) at the ciliary tip in WT and *BBS1* KO cells using a fluorescence recovery after photobleaching (FRAP) approach (Fig. 4C-D). Since IFT trains pause and remodel at the ciliary tip before engaging retrograde motors and cargoes, measuring their dwell time (halftime of recovery) serves as an indirect readout of retrograde transport initiation. In BBSome-deficient cells, IFT74–mNG fluorescence at the ciliary tip recovered significantly faster (half-time, T□/□ ≈ 9.1 s) compared with WT cells (T□/□ ≈ 15.3 s) (Fig. 4E-F). The shorter dwell time of IFT in the absence of the BBSome likely reflects a failure to engage BBSome-dependent cargoes, resulting in earlier initiation of retrograde transport. Introduction of the BBS1 WT variant into IFT74–mNG *BBS1* KO cells (Fig. 4D, G) restored IFT dwell time at the ciliary tip to comparable value (TL/L ≈ 13.2 s) observed in WT cells (T□/□ ≈ 15.3 s) (Fig. 4F, H). In contrast, IFT dwell time at the ciliary tip decreased in cells expressing the BBS1 R160Q–containing BBSome (T□/□ ≈ 6.8 s), phenocopying the short dwell time observed in BBSome-deficient cells (T□/□ ≈ 9.1 s) (Fig. 4F, H). These data indicate that the BBS1 R160Q variant prevents IFT to load the TOM1L2-dependent cargoes, resulting in earlier departure of retrograde IFT trains.

Collectively, our results demonstrate that BBS1 mutations linked to BBS fall into distinct functional classes defined by their impact on BBSome assembly and ciliary function (Fig. 5). M390R and E224K disrupt BBSome assembly at different stages, impairing pre-BBSome formation and BBSome maturation/ciliary translocation, respectively. In contrast, R160Q permits BBSome assembly and ciliary localization but selectively disrupts BBSome-dependent TOM1L2-mediated cargo export via IFT.

**Figure 5.**
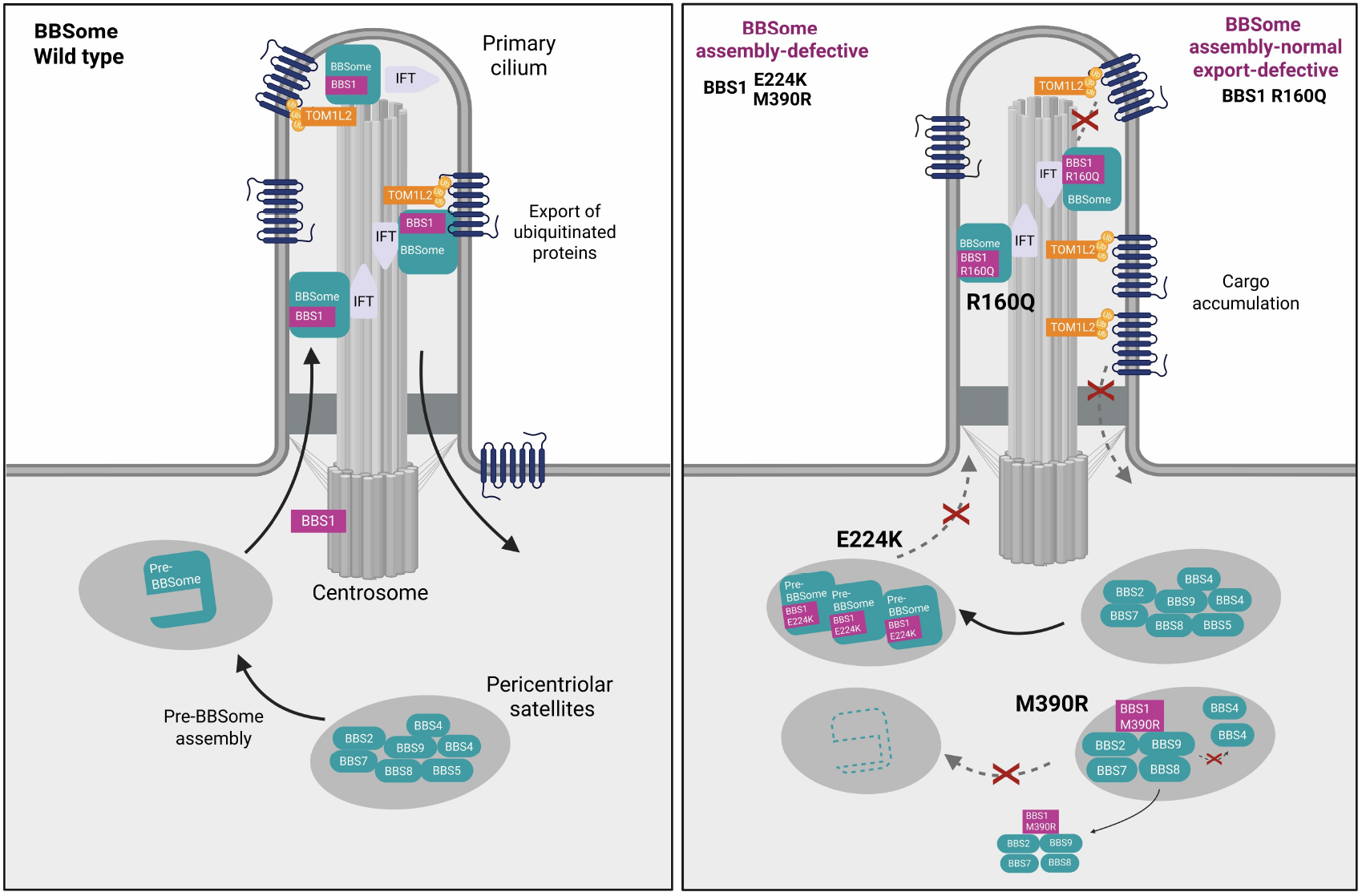
BBS1 mutation–specific defects in BBSome assembly and function define distinct pathogenic classes. In WT condition, BBS4 at pericentriolar satellites nucleates pre-BBSome assembly, which is completed by BBS1 to form a stable, mature BBSome that enters the cilium. Within cilia, the BBSome cooperates with IFT to mediate export of TOM1L2-marked, ubiquitinated cargoes. BBS1 mutations linked to Bardet–Biedl syndrome can be divided into two functional classes based on how they impair BBSome assembly and function in cilia. M390R and E224K belong to a class of mutations that disrupt distinct stages of BBSome assembly and maturation: M390R impairs initial pre-BBSome assembly at satellites, whereas E224K disrupts BBSome maturation and prevents its translocation from satellites to the cilium. In contrast, R160Q represents a second class of mutations that supports BBSome assembly and ciliary localization but impairs BBSome function within cilia. Specifically, R160Q prevents TOM1L2-marked cargo loading onto IFT, resulting in defective GPCR export and increased IFT turnover.

## Discussion

Bardet–Biedl syndrome (BBS) is a ciliopathy characterized by striking clinical heterogeneity, even among individuals carrying mutations in the same gene ^4,22,28,29,42^. Mutations in BBS1 are distributed throughout the gene without clear mutational hotspots, and missense variants generally cause milder disease than loss-of-function mutations (Beales et al., 2003; Chou et al., 2019; Niederlova et al., 2019). In this study, we investigated the molecular mechanisms underlying the pathogenicity of three BBS1 missense variants - M390R, E224K, and R160Q – displaying distinct levels of clinical penetrance ^4,27–31^.

The M390R mutation is the most prevalent variant among BBS patients (∼80% of BBS cases) ^4,22^. Our data, together with previous studies, provide a mechanistic explanation for the pathogenic effects of the M390R mutation, demonstrating that it destabilizes the BBS1 β-propeller structure ^23,26^, and weakens interactions with the BBS1 partners BBS4 and BBS5. These combined defects impair productive pre-BBSome assembly at pericentriolar satellites ^5,33^. Notably, the association of the BBS1 M390R variant with the BBSome core (BBS2/7/9) indicates that this mutation does not disrupt BBS chaperonin-mediated stabilisation of BBS2 and BBS7 ^32,43^. Instead, the BBS1 M30R variant prevents recruitment of the BBSome core to BBS4 and sequesters the core complex from satellites into the cytoplasm. At the cellular level, the M390R mutation approximates a functional BBSome-null state, explaining its broad clinical spectrum despite BBS1 being considered a “milder” BBS gene ^4^. In addition, the M390R variant may exert further pathogenic effects through altered interactions with BBS4, a core satellite-associated protein ^5,44^. Dysregulation of BBS4 dynamics could disrupt satellite composition and homeostasis, impairing centrosomal protein trafficking, ciliogenesis, and the cell-cycle-related functions ^44–48^. Such defects likely exacerbate BBSome deficiency and underlie the severe BBS1 M390R phenotype.

In contrast, the E224K mutation represents an intermediate defect, producing sub-threshold BBS features that can lead to misdiagnosis as Alström syndrome or to an unconfirmed diagnosis due to limited evidence supporting its pathogenicity ^29,34^. The BBS1 E224K variant permits pre-BBSome assembly at satellites and associates with this complex; however, the resulting BBSome–BBS1 E224K fails to translocate to the cilium, likely via a BBS3-dependent mechanism ^6,25,49^. Our data and structural analyses point to incomplete complex maturation, reflecting weakened integrity that may be critical for complex stability and activation ^7,25,26,49^.

The R160Q substitution is a result of 5` splice site mutation compromising splicing efficiency and expression of BBS1 ^31,50^. The R160Q variant has been linked to milder phenotypes or, in some cases, isolated non-syndromic retinitis pigmentosa ^28–31^. The R160Q elevates ciliary TOM1L2, indicating impaired BBSome–TOM1L2 recognition, a key step in BBSome-dependent retrieval of ubiquitinated cargoes from cilia ^10,11,19^. The milder penetrance of R160Q mutation is intriguing, as both E224K and M390R also increase ciliary TOM1L2. One possible explanation is that R160Q selectively impairs BBSome-dependent cargo export while preserving BBSome localisation. Moreover, TOM1L2-dependent retrieval may be particularly critical in photoreceptors, which rely on continuous protein and lipid turnover to maintain membrane homeostasis ^18,19,51,52^, providing a rationale for the predominantly retinal phenotype associated with R160Q mutation.

To enter the cilium, the BBSome is loaded onto anterograde IFT trains at the ciliary base, functioning both as a cargo for IFT and as an adaptor for BBSome-dependent transmembrane ciliary proteins ^6,9,38,53,54^. There is limited knowledge about how the BBSome regulates IFT dynamics in mammalian cilia, despite clear evidence supporting their genetic interaction in BBS patients and mouse models ^12,55^. In our study, we leveraged the BBS1 R160Q variant to provide the first insights into IFT behavior directly within cilia in the context of a BBS-linked mutated BBSome. Since the BBSome-BBS1 R160Q reaches the cilium, its association with the anterograde IFT trains is most likely preserved ^9,53^. Upon IFT train remodeling at the ciliary tip, retrograde IFT-BBSome-BBS1 R160Q trains fail to efficiently load TOM1L2-marked cargo, resulting in accelerated or premature retrograde IFT. This kinetic defect is most likely compatible with normal ciliogenesis and basal ciliary architecture but may impair dynamic signaling responses, consistent with the subtle metabolic and retinal phenotypes observed in milder BBS cases.

Finally, it remains to be determined whether cilia-localised BBSome-BBS1 R160Q is sufficient to prevent other extraciliary defects associated with BBSome deficiency, including disrupted planar cell polarity and WNT signalling, increased extracellular vesicle release, mitochondrial dysfunction, and altered cell cycle progression ^15,44,56,57^, which will define how BBSome dysfunction drives BBS pathology of varying severity.

In summary, our study reveals that BBS1 mutations disrupt BBSome function through distinct mechanisms that align with clinical severity. Defects in BBSome assembly and trafficking correlate with more penetrant disease, whereas preserved assembly with selective export failure underlies milder phenotypes. Our findings provide a mechanistic bridge between genotype and phenotype in BBS and highlight the importance of dissecting mutation-specific BBSome dysfunction to inform diagnosis, prognosis, and therapeutic development.

## Methods

### Antibodies, dyes and reagents

Mouse anti-acetylated tubulin antibody (IF, 1:50) was provided by Dr. Vladimir Varga, Institute of Molecular Genetics of the Czech Academy of Sciences. Mouse anti-FLAG (M2 clone) (F1804; WB, 1:1000), mouse anti-β-Actin (A1978, WB, 1:5000), rabbit anti-BBS9 (HPA021289; IF, 1:250, WB, 1:500) and rabbit anti-BBS7 (HPA044592; WB, 1:500) were purchased from Sigma Aldrich. Mouse anti-PCM-1 (sc-398365; IF, 1:50), mouse anti-BBS2 (sc-365355; WB, 1:500) and mouse anti-BBS8 (sc-271009; WB, 1:500) were purchased from Santa Cruz. Rabbit anti-BBS1 (ab166613; WB, 1:500) and rabbit anti-TOM1L2 (ab96320; IF, 1:200) were purchased from Abcam. Rabbit anti-GPR161 (13398-1-AP; IF, 1:200), rabbit anti-BBS5 (14569-1-AP; WB, 1:500) and rabbit anti-BBS4 (12766-1-AP; WB, 1:500) were purchased from Proteintech. Rabbit anti-mCherry (PA534974; IF, 1:100) was purchased from ThermoFisher. Goat anti-GFP (ab6673; IP, 3.2 µg per sample) was purchased from Abcam. Primary conjugated antibody for flow cytometry anti-CD90.1 PE-Cy7 (25-0900) was purchased from eBioscience.

Secondary antibodies for IF were as follows: anti-mouse Alexa Fluor 647 (Invitrogen, A-21235; 1:1000), anti-rabbit Alexa Fluor 488 (Invitrogen, A11008; 1:1000). Secondary antibodies for WB were as follows: goat anti-rabbit-HRP (Jackson, 111-035-144; 1:10,000), goat anti-mouse-HRP (Jackson, 115-035-146; 1:10,000), bovine anti-goat-HRP (Jackson, 805-035-180; 1:10,000).

SiR-Tubulin (SC002) was purchased from Spirochrome and used at a final concentration of 1 µM. ProLong™ Gold antifade reagent with 4′,6-diamidino-2-phenylindole (DAPI) (P36941) was purchased from Invitrogen.

### Cell cultures and treatments

Immortalized human retinal pigment epithelium cell line, hTERT-RPE-1 (RPE1) (ATCC, CRL-400) was provided by Dr. Vladimir Varga (Institute of Molecular Genetics of the Czech Academy of Sciences). RPE1 *BBS1* KO cell line was established previously ^5^. Phoenix Ampho cell line was provided by Dr. Tomas Brdicka (Institute of Molecular Genetics of the Czech Academy of Sciences All cell lines were cultured in complete Dulbecco’s modified Eagle’s medium (DMEM, Sigma, D6429 - 500 mL) supplemented with 10% fetal bovine serum (FBS), 100 U/mL penicillin (BB Pharma), 100 μg/mL streptomycin (Sigma-Aldrich), and 40 μg/mL gentamicin (Sandoz). All cell lines are regularly tested for mycoplasma contamination.

Smoothened agonist - SAG (566660; Sigma-Aldrich) was used at a concentration of 200 nM. The treatments were for 2 h if not indicated otherwise.

### Cloning and gene transfections

*BBS1* ORF was amplified from pMSCV-BBS1-YFP ^5^, appended with mCherry and FLAG coding sequences at the N terminus using recombinant PCR, and cloned pMSCV-IRES-Thy 1.1 vector (Clontech) using BglII and EcoRI restriction sites. PCR based mutagenesis was performed on the mCherry-FLAG-BBS1 open reading frame (ORF), where selected BBS1 mutations were introduced using designed primers: M390R (c.1169T>G, F: CACCCTCATCAGGACCACTCGA, R: TCGAGTGGTCCTGATGAGGGTG), E224K (c.670G>A, F: TGCTGGGCACCAAGAACAAGGA, R: TCCTTGTTCTTGGTGCCCAGCA) and R160Q (c.479G>A, F: GAGAGCATCCAGGAGACGGCAG, R: CTGCCGTCTCCTGGATGCTCTC).

The viral transduction of cell lines was done according to ^58^ with slight modifications. For viral particle production, Phoenix Ampho cells were seeded on 10 cm dish and allowed to reach confluency of 60-70%. 30 µg of plasmid DNA was transfected into Phoenix Ampho cells using the polyethyleneimine to generate retroviruses. Cells were incubated overnight, and the media was changed to production media (DMEM+ATB+FBS) on the next day. RPE1 cells were transduced with 2 mL of supernatant containing viral particles along with 8 µg/mL polybrene and analyzed for transgene expression after 48 h. RPE1 cells stably expressing both mCherry and Thy 1.1. markers or YFP were bulk sorted using FACS Aria IIu (BD Biosciences).

### Protein lysates, co-immunoprecipitation and western blotting

RPE1 cells were grown in 15 cm cell culture dishes and lysed in the lysis buffer (20mM HEPES at pH 7.5, 150mM NaCl, 2mM EDTA at pH 8, 0.5% Triton X-100) supplemented with protease inhibitor cocktail (complete, Roche, #05056489001). Lysates were cleared by centrifugation at 15,000 × g for 15 minutes at 4°C. Protein concentration was adjusted using the Pierce™ BCA Protein Assay Kit (Thermo Scientific). For co-immunoprecipitation assay, protein lysates were immunoprecipitated with anti-FLAG M2 beads (Sigma Aldrich, A2220) or goat anti-GFP antibody (Abcam, 6673, 3.2 µg per sample) conjugated to Protein A Plus UltraLink beads (ThermoFisher, #53142) for 1.5 h at 4°C. Beads were washed three times in lysis buffer and co-precipitated proteins were denatured in 1× Laemmli buffer. For protein expression analysis, protein lysates were immediately denatured in reducing 4× Laemmli buffer. Denatured protein samples were analyzed by Western blotting following standard protocols. Membranes were probed with primary antibodies overnight at 4°C and secondary antibodies for 1 h at room temperature. Membranes were developed using chemiluminescence immunoblot imaging system Fusion Solo S (Vilber).

### Flow cytometry

RPE1 cells expressing the mCherry-FLAG-tagged BBS1 variants were cultured in 6-well dishes, trypsinized and centrifuged at 400 × g for 4 minutes. The cell pellets were washed in PBS and resuspended in FACS buffer (2 mM EDTA, 2% FBS, and 0.1% sodium azide) containing anti-CD90.1 PE-Cy7 antibody (eBioscience, 25-0900). Measurements were taken on Cytek Aurora Flow Cytometer (Cytek). Geometric mean fluorescence intensities of mCherry and Thy 1.1 positive cells were obtained and used to examine the protein expression. Flow cytometry data was analyzed in FlowJo software, version 10.x (BD Life Sciences).

### Immunofluorescence

RPE1 cells were seeded on 12 mm coverslips and serum starved for 24 h. After respective treatments, cells were fixed (4% formaldehyde) and permeabilized (0.2% Triton X-100) for 10 min. Optional step after fixation was antigen retrieval (BBS9 - Fig. 2A, TOM1L2 - Fig. 4A), when cells were incubated for 5 minutes in 95°C in the pre-heated antigen retrieval buffer (100 mM TRIS, 5% urea, pH 9.5). Blocking was done using 5% goat serum (Sigma, G6767-100 mL) in PBS for 15 min and incubated with primary antibody (1% goat serum/PBS) and secondary antibody (PBS) for 1 h and 45 min, respectively in a wet chamber. The cells were washed after each step in PBS three times. At last, the cells were washed in distilled H_2_O, air-dried, and mounted using ProLong™ Gold antifade reagent with DAPI (P36941; Invitrogen).

### Fluorescence microscopy and data analysis

Image acquisition for BBS9, mCherry and GPR161 localization was performed on the Delta Vision Core microscope using the oil immersion objective (Plan-Apochromat 60× NA 1.42) and filters for DAPI (435/48), FITC (523/36), TRITC (576/89) and Cy5 (632/22). Z-stacks were acquired at 1024 × 1024-pixel format and Z-steps of 0.2 µm.

Image acquisition for TOM1L2 localization was performed on the Leica DM6000 microscope using the oil immersion objective (HCX PL APO 63× NA 1.40) and filter cubes for DAPI (LP 425), GFP (525/50), TRITC (600/40) and Cy5 (700/75). Z-stacks were acquired at 2048 × 2048-pixel format and Z-steps of 0.2 µm.

Maximum intensity projections were used to quantify the frequency of BBS9 localization to cilia (Fig. 2B), BBS9 localization to PS (Fig. 3B), and TOM1L2 localization to cilia (Fig. 4B), as well as to measure BBS9 intensity profiles (Fig. 2C) and GPR161 signal intensity (Fig. 1C) within cilia using the Fiji ImageJ. Signal intensity was quantified using the line tool following subtraction of background signal adjacent to the cilium. Intensity distribution along the cilium was analysed using the Plot Profile function with background subtraction, followed by normalization to the first data point to facilitate comparison between conditions.

### Fluorescence recovery after photobleaching, data processing, and analysis

Cells expressing YFP-BBS4 or IFT74-mNeonGreen were seeded into glass-bottom 8-well chambers (CellVis, USA) and serum-starved for 24 h or 48 h, respectively. On the day of the FRAP experiments, cells were washed and incubated for at least 1 h at 37°C with 5% CO_2_ in FluoroBrite medium supplemented with 1μM SiR-Tubulin (Spirochrome, SC002) to visualize centrosomes or cilia. Prior to live-cell imaging, 20mM HEPES was added to the medium. FRAP measurements were performed using a Leica TCS SP8 confocal microscope equipped with an oil immersion objective HC

Plan-Apochromat 63× NA 1.4 oil, CS2 at room temperature. Data acquisition was performed in 512×512-pixel format with pinhole 2.62 Airy, at a speed of 1000 Hz (BBS4) or 400 Hz (IFT74) in bidirectional mode and 8-bits resolution. Photobleaching of satellites (0.3 s) was performed with a circular spot (1 µm in diameter), and of ciliary tip (0.65 s) with a rectangle roi (∼ 1 µm x 1 µm) at 100% intensity using a 20 mW 488 nm solid-state laser. Fluorescence recovery was monitored at low laser intensity (3-10%) at 0.26 s (satellites) or 0.648 s (ciliary tip) intervals until reaching the plateau of recovery, in total for 60 to 80 s after photobleaching. A total of 30-50 separate FRAP measurements were performed for each sample. All FRAP curves were normalized to fluorescence loss during acquisition following the subtraction of background fluorescence. Curve fitting was performed in GraphPad Prism software using the one-phase association fit. All individual curves were fitted at once to obtain the mean and 90% or 95% confidence intervals of the desired parameters, halftime T_1/2_ and the mobile fraction F_m_. Since the initial recovery after bleaching is contaminated by the diffusion, for simplicity and to estimate only the halftime of the protein fraction bound at the analyzed compartments, we restricted fitting of our data to times t > 2 s as we did in our previous studies ^5,59^.

### Statistical analysis

Statistical analysis was performed using GraphPad Prism Version 5.04. The following statistical tests were used and are indicated in the respective Figure legends: Mann-Whitney test, Paired t-test and Spearman correlation.

## Data availability

This study includes no data deposited in external repositories.

## Acknowledgement

This project has received funding from the Czech Science Foundation (24-12431S to MH, 25-17600S to DR), core funding provided by the Institute of Molecular Genetics of the Czech Academy of Sciences (RVO 68378050). We acknowledge the Light Microscopy Core Facility, IMG, Prague, Czech Republic, supported by MEYS – LM2023050 and RVO – 68378050-KAV-NPUI, for their support with the widefield and confocal imaging and data analysis presented herein.

## Author contribution

MH conceived the study and was in charge of the overall direction and planning. KM and AP performed cloning and cell line generation.KM performed most of the basic fluorescence microscopy, biochemical experiments and data analysis. HH, SB, SS and MH performed basic and quantitative fluorescence microscopy and data analysis. DR prepared the structural model. MH wrote the manuscript with the contribution of all the other authors.

## Additional Declarations

The authors have declared there is **NO** conflict of interest to disclose

