## Supplemental Information for "Bardet-Biedl Syndrome 1 Mutations Differentially Impact BBSome Integrity and its Function in Ciliary Trafficking"

**Supplemental Figure 1. BBS1 variants are stable and equally expressed.**

(A) Gating strategy for flow cytometric analysis of the mCherry-FLAG tagged BBS1 variants expressing *BBS1* KO RPE1 cell lines. Debris was excluded using FSC-H/SSC-H, followed by singlet discrimination using FSC-A/FSC-H and SSC-H/SSC-A. Marker-positive populations were defined using fluorescence-minus-one controls and used for downstream quantification.

(B) Correlation plot of the expression levels (gMFI) of the mCherry-FLAG tagged BBS1 variants and Thy1.1 expressed from the same retroviral constructs in *BBS1* KO RPE1 cell lines. The cell line populations were divided into 5 subsets based on the Thy1.1 signal as shown in (A). Linear regression is indicated by the solid line, with the Spearman correlation coefficient r (very strong correlation 0.80 – 1.00) and p value for each condition. Representative plot out of three independent experiments is shown.

(C) Expression levels of endogenous and exogenous BBS1 subunit in parental and BBS1 variants expressing WT and *BBS1* KO RPE1 cell lines. Equal protein amounts were loaded into each lane. Actin is used as the loading control. Representative blots out of three independent experiments are shown.

Bar graphs depicting the relative expression levels of the BBS1 subunit (D) and endogenous BBSome subunits (E) in parental and BBS1 variants expressing WT and *BBS1* KO RPE1 cell lines. Protein amounts were quantified using the Fiji ImageJ. Protein expression was first normalized to actin and then to levels in parental WT cell line. Average and SD of three independent experiments is shown.

**Supplemental Figure 2. Localization of the mCherry-FLAG tagged BBS1 variants**

(A) Representative micrographs depicting localisation of mCherry-FLAG BBS1 variants in the WT and *BBS1* KO RPE1 cell lines upon 24 h serum starvation. Antibodies against mCherry and acetylated tubulin (Ac-tub) were used to visualise BBS1 and the primary cilia respectively. Nucleus was stained with 4′,6-diamidino-2-phenylindole (DAPI). Scale bar, 5µm.

(B) Quantification of cilia frequency in the parental and BBS1 variants expressing WT and *BBS1* KO RPE1 cell lines upon 24 h serum starvation assessed in Fig. 2A. Analysis was carried out using the Fiji ImageJ. Mean and SD from three independent experiments (n = 330-620 cells).

(C) Quantification of the co-precipitated BBSome subunit BBS8 in Fig. 2D was performed using the Fiji ImageJ. Amount of the BBS8 was normalized to the levels of the mCherry-FLAG tagged BBS1 WT in *BBS1* KO cell line detected on the respective membrane. Mean with SD of three experiments is shown.

**Supplemental Figure 3. YFP-BBS4 dynamics at pericentriolar satellites in the presence of BBS1 variants**

(A) Representative micrographs depicting localisation of YFP-BBS4 in the parental and BBS1 variants expressing WT and *BBS1* KO RPE1 cell lines. Centrosome and microtubules were stained with 1µM SiR-tubulin. Scale bar, 5µm.

(B) FRAP analysis of the dynamic turnover of the YFP-BBS4 in the presence of BBS1 WT and E224K variants at the pericentriolar satellites in *BBS1* KO RPE1 cell lines upon 24 h serum starvation. Recovery curves were fitted at once using one phase association fit. Mean of 34 (WT) and 29 (E224K) measurements from four independent experiments is shown.

Bar graphs depicting recovery halftimes (s) (C) and mobile fractions (D) of YFP-BBS4 in the presence of BBS1 WT and E224K variants at the pericentriolar satellites in *BBS1* KO RPE1 cell lines obtained from FRAP analysis in B. Means of 34 (WT) and 29 (E224K) measurements from four independent experiments are shown. Error bars represent the 90% confidence interval and p-value is shown.

(E) Co-immunoprecipitation of the BBSome subunits from YFP-BBS4 expressing WT and *BBS1* KO RPE1 cell lines using anti-GFP antibodies after 24 h serum starvation. Representative immunoblot out of three experiments is shown.

(F) Quantification of the co-precipitated BBSome subunits in (E) was performed using the Fiji ImageJ. Amount of the BBSome subunits was normalized to the YFP-BBS4 levels in WT RPE1 cell line, as detected on the respective membrane. Mean with SD of three experiments is shown.

Supplemental Figure 1

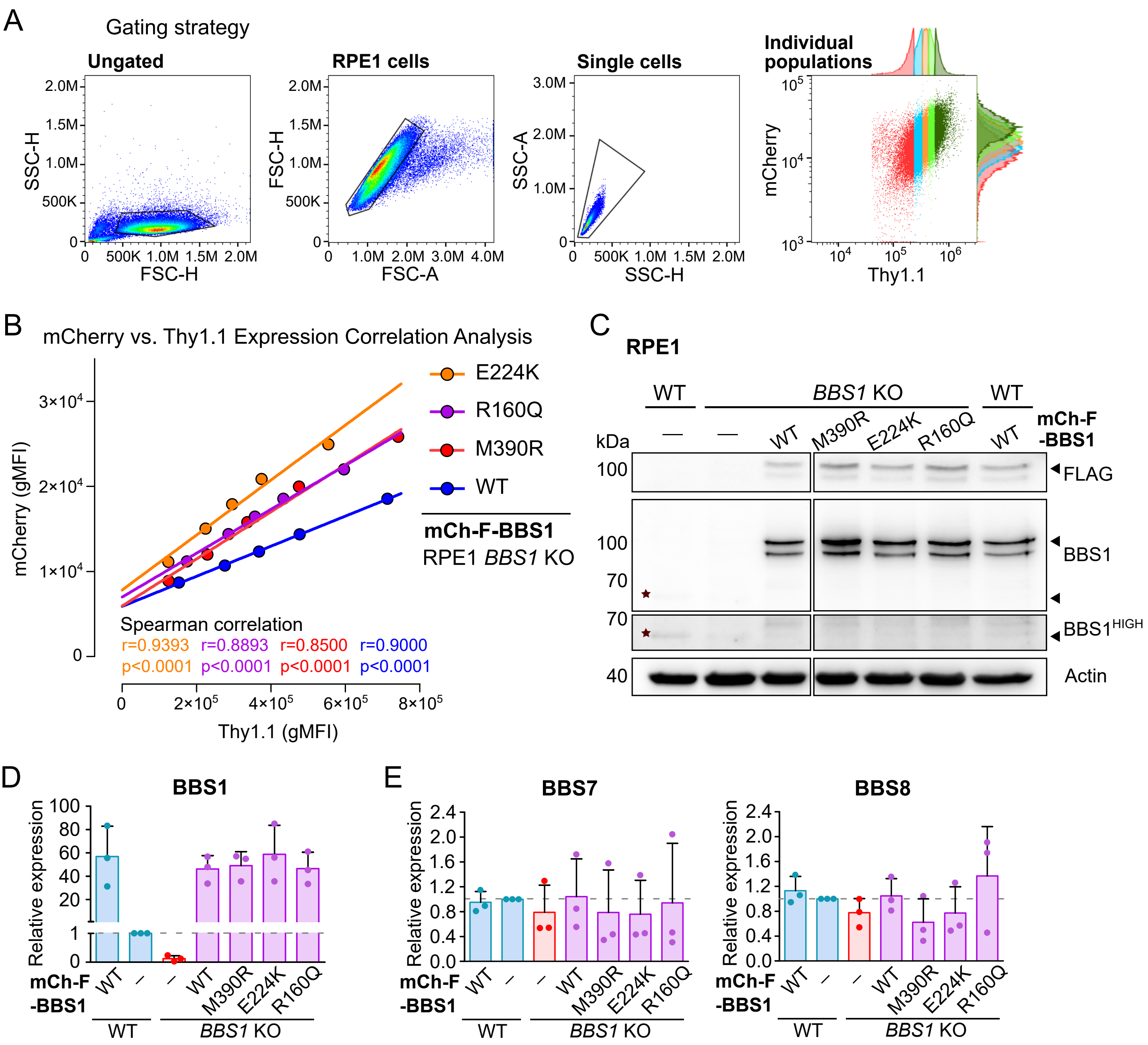

Supplemental Figure 2

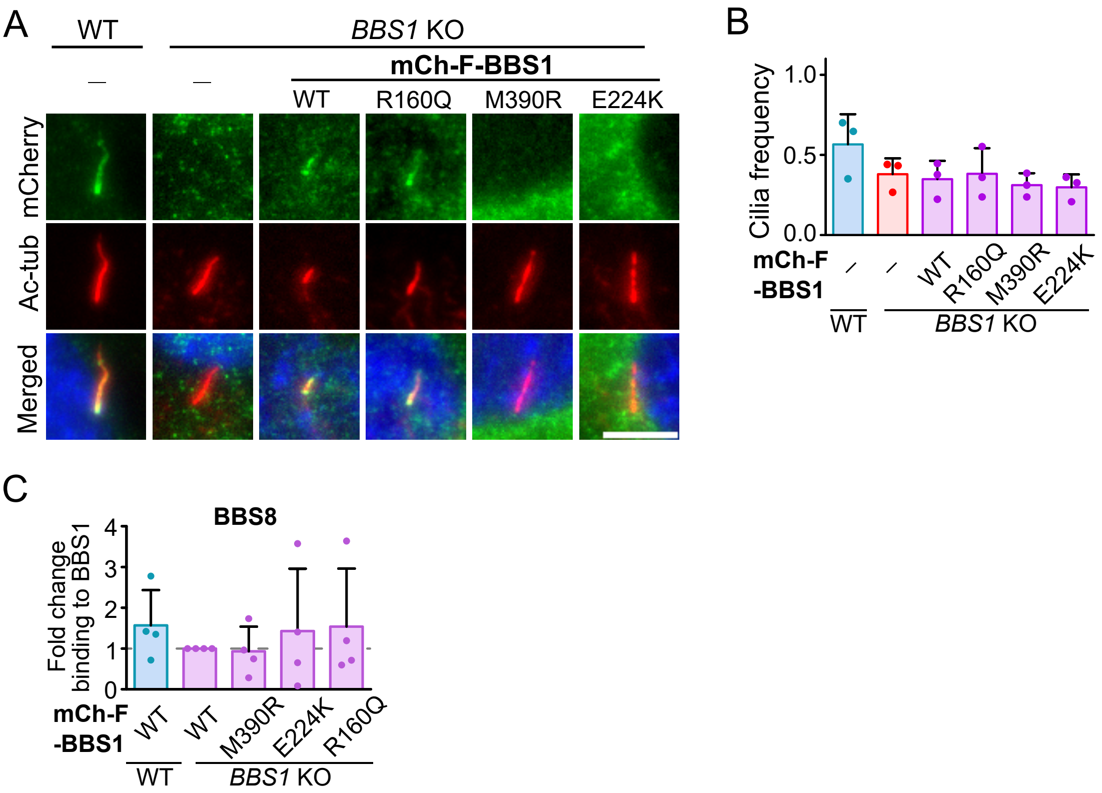

### Supplemental Figure 3

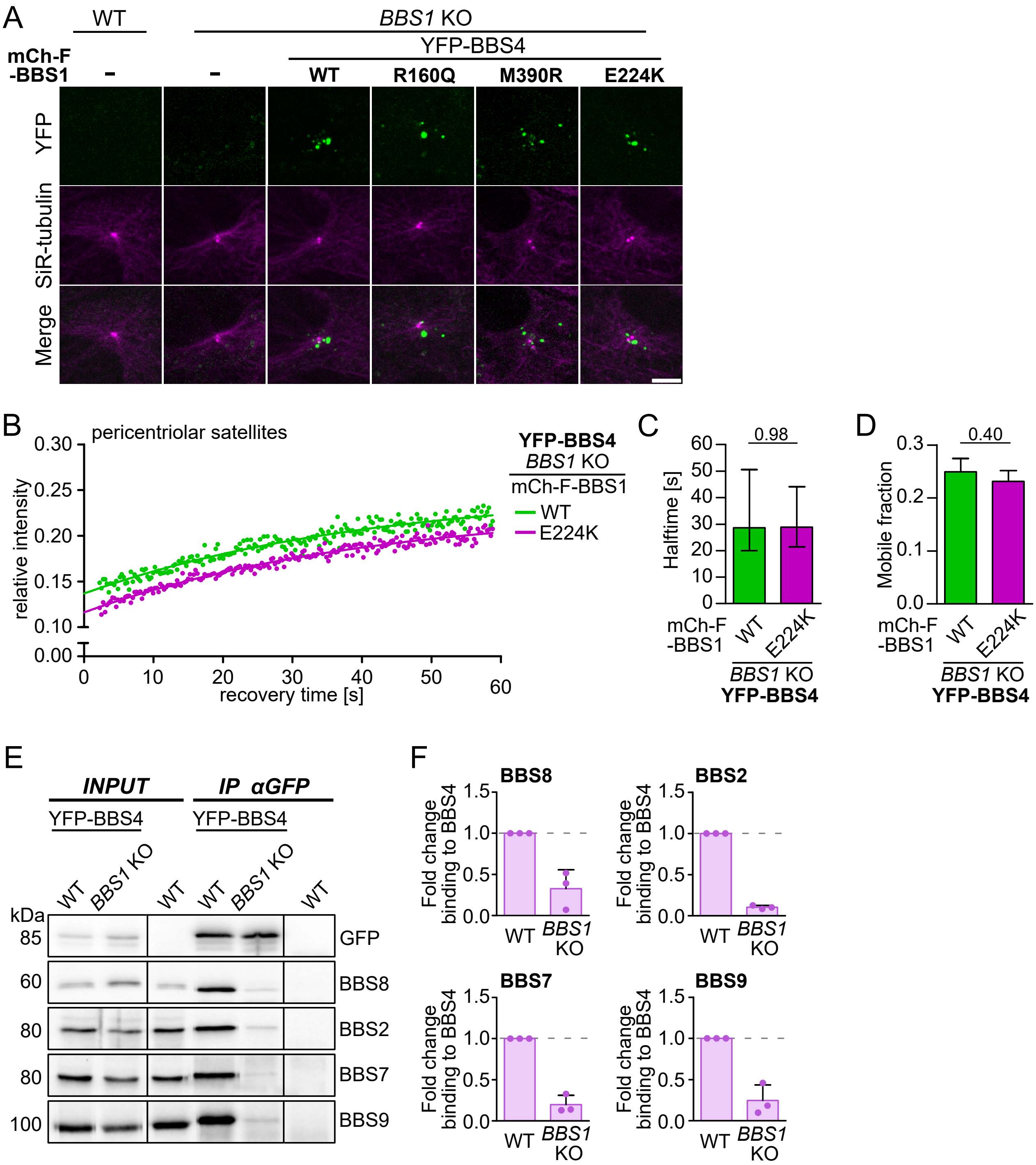
